# Compendium of Synovial Signatures Identifies Pathologic Characteristics for Predicting Treatment Response in Rheumatoid Arthritis

**DOI:** 10.1101/279786

**Authors:** Ki-Jo Kim, Minseung Kim, Ilias Tagkopoulos

## Abstract

Treatment of patients with rheumatoid arthritis (RA) is challenging due to clinical heterogeneity and variability. Integration of RA synovial genome-scale transcriptomic profiling of different patient cohorts can provide insights on the causal basis of drug responses. A normalized compendium was built that consists of 256 RA synovial samples that cover an intersection of 11,769 genes from 11 datasets. Differentially expression genes (DEGs) that were identified in three independent methods were fed into functional network analysis, with subsequent grouping of the samples based on a non-negative matrix factorization method. Finally, we built a predictive model for treatment response by using RA-relevant pathway activation scores and four machine learning classification techniques. We identified 876 up-regulated DEGs including 24 known genetic risk factors and 8 drug targets. DEG-based subgrouping revealed 3 distinct RA patient clusters with distinct activity signatures for RA-relevant pathways. In the case of infliximab, we constructed a classifier of drug response that was highly accurate with an AUC/AUPR of 0.92/0.86. The most informative pathways in achieving this performance were the NFκB-, FcεRI- TCR-, and TNF signaling pathways. Similarly, the expression of the HMMR, PRPF4B, EVI2A, RAB27A, MALT1, SNX6, and IFIH1 genes contributed in predicting the patient outcome. Construction and analysis of normalized synovial transcriptomic compendia can provide useful insights for understanding RA-related pathway involvement and drug responses for individual patients. The efficacy of a predictive model for personalized drug response has been demonstrated and can be generalized to several drugs, co-morbidities, and other relevant features.

## Introduction

Rheumatoid arthritis (RA) is a complex autoimmune disease involving a multitude of environmental and genetic factors that exhibit nonlinear dynamic interactions (1). The disease is characterized by chronic inflammation of the synovium, which results in irreversible damage to the joints over time, leading to pain and functional impairment. Severity and clinical course of the disease is highly variable across the different patients and hence difficult to predict (1). Despite an introduction of tumor necrosis factor (TNF) inhibitors, over 30% of patients do not respond fully to therapy (2). Moreover, a considerable subset of the patients who showed initial good response experience disease flare or efficacy reduction even on drugs (2). A similarly discouraging picture is painted in the case of other biologics, which creates a current and present need for better understanding of the disease. An early, aggressive and personalized treatment that provides the best possible drug combination for a patient is likely to improve our ability to treat RA. Despite the introduction of novel drugs and the fact that RA is an active research topic, however, we still have substantial gaps in our knowledge regarding the mechanistic basis of RA progression, which is needed to administer personalized and precise care.

In RA, gene expression profiling has been used to gain insights regarding pathogenesis and drug response (3). Since studies so far have been in unrelated cohorts and study groups, small sample size, heterogeneity in study population (sex, age, and ethnicity), differences in technical protocols, microarray platform, and data analysis methods has hindered a comprehensive analysis across all available datasets. In addition, most studies have collected samples from whole blood or peripheral blood mononuclear cells, which are easier to acquire but have a limited capacity to adequately reflect the joint inflammation (4-6). However, integrated analysis of the compendium by the accrued genome-wide datasets provided opportunities to capture the missing features, bridge the gap between prior knowledge, and better understand human diseases in the field of cancer and infectious disease (7,8).

In this study, our aim is to elucidate the various transcriptional and signaling signatures of RA by performing a comprehensive meta-analysis of the publicly available datasets that have been published so far. We focus on synovial tissue samples to avoid the high false discovery rates coming from blood samples. We have applied several preprocessing and normalization steps to create a cohesive, homogenized compendium of genome-wide gene expression signatures for downstream analysis. We used this compendium to separate expression-driven subgroup, understand the key cellular components in each group and then use genes and pathways with high information value that we have identified to create predictive models for drug responsiveness.

## Methods

### Systematic search and data collection

We used the keywords “Rheumatoid Arthritis”, “Synovium or synovial tissue”, “Transcriptomics or microarray”, “Dataset” in Google Scholar and PubMed to find relevant publications to the topic of synovial gene signatures of patients with rheumatoid arthritis (**Fig. 1**). We retrieved all publications that were accompanied by high-throughput datasets (20 studies in total). From the resulting set, we removed entries that had been duplicated and selected datasets measuring over 10,000 genes to secure the largest size of genes and samples. Since there was a trade-off between the number of studies to include and the number of genes that are within the intersection from all datasets, we optimized the product of the two by selecting the point where these two trends cross (**Fig. S1**). The final RA sample count was 256, the osteoarthritis (OA) count 41, and 36 normal (NC) samples were included as controls. Ultimately, the final RA compendium was constructed out of 11 studies with a total of 333 samples, one per patient, covering 22,721 genes total (common core of 11,769 genes).

**Figure 1.**
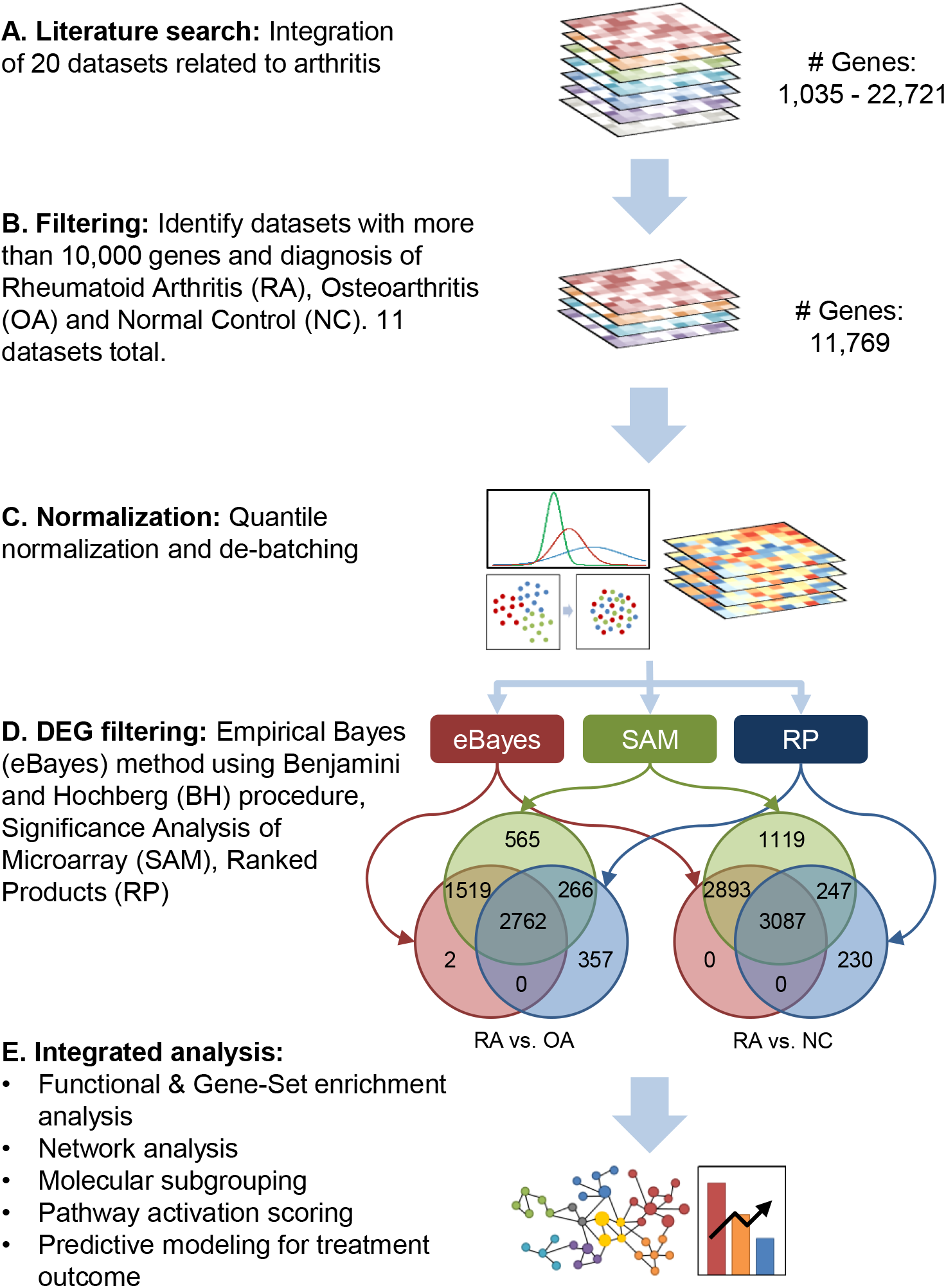
Overview of the data processing steps. **(A)** Twenty studies maximally covering 20,511 genes were retrieved from the literature. **(B)** Selected were 11 datasets adequate to integrated analysis, which included 256 RA, 41 OA, and 36 NC samples covering 11,769 gene. **(C)** The merged dataset was normalized using quantile method and its batch effect was corrected. **(D)** DEG of RA compared to OA or NC were obtained using three methods, eBayes, SAM, and RP. Intersection of three DEG sets was chosen as significant DEG. The number of DEG was 2762 in RA versus OA and 3087 in RA versus NC. (E) A list of strategies for integrated analysis. (Abbreviation: RA, rheumatoid arthritis; OA, osteoarthritis; NC, normal controls; DEG, differentially expressed genes; eBayes, empirical Bayes; SAM, significance analysis of microarray; RP, rank products).

### Data normalization and removal of batch effects

For one-channel arrays, the image data was first imported and then the Robust Multi-array Average (RMA) method was applied for a set of replicates for background correction, normalization, probe-set summarization. For dual-channel arrays, the image data were imported and background correction was performed using normexp as it was shown to outperform other methods. Red and green channels were separated and quantile-normalized for each set of replicates. The vectors for the matrices were normalized using the quantile normalization method. Residual technical batch effects arising due to heterogeneous data integration were corrected using the ComBat function within the empirical Bayes package. Quality assurance and distribution bias was evaluated by Principal Component Analysis (**Fig. S2**).

### The RA compendium

After preprocessing, the gene expression profiles have a significant reduction of systematic, dataset-specific bias in comparison with the same dataset before normalization and batch correction (**Fig. S2**). The resulting compendium has a gene size of 11,769 in 333 samples, including 256 RA patients, 41 OA patients, and 36 normal controls. In 105 of the RA samples, synovial tissue sampling was conducted before the start of certain drug: 11 for adalimumab (ADLM), 62 for infliximab (IFXM), 8 for methotrexate (MTX), 12 for rituximab (RTXM), and 12 for tocilizumab (TOCM). For these patients, assessment of disease activity and response was performed per the EULAR response criteria (9) 12-16 weeks after initiation of therapy: 32 were good, 47 were moderate, and 26 were poor responders. Information on demographics and clinical characteristics including age, sex, disease duration, and RF positive were not fully annotated for each RA sample (**Table S1**).

### Filtering of differentially expressed genes

In order to identify the differentially expressed genes (DEGs), we employed three widely-used methods: (a) an empirical Bayesian method using the Benjamini and Hochberg procedure with a significance threshold at an adjusted p-value < 0.05; (b) the Significance Analysis of Microarray (SAM) method, with a significance threshold of false discovery rate FDR < 0.05; (c) the Rank Products (RP) method with a significance threshold set at percentage of false prediction pfp < 0.05. The resulting list of DEGs is the intersection of the three individual DEGs sets for each method to minimize the false discovery rate statistic.

### Functional enrichment analysis

We performed functional enrichment analysis focusing on the up-regulated DEGs using the Database for Annotation, Visualization, and Integrated Discovery (DAVID) software (10). Terms were regarded significant if the p-value (EASE score) is lower than 0.05, the enrichment score higher than 1.3, and the fold enrichment was larger than 1.5.

### Gene set enrichment analysis

Gene set enrichment analysis (GSEA) analysis was carried out using the GSEA software from the Broad Institute to assess the overrepresentation of RA-related gene sets (11). The enrichment results were visualized with the Enrichment Map format, where nodes represent gene-sets and weighted links between the nodes represent an overlap score depending on the number of genes two gene-sets share (Jaccard coefficient) (12). To intuitively identify redundancies between gene sets, the nodes were connected if their contents overlap by more than 25%. Clusters map to one or more functionally enriched groups, which were manually circled and assigned a label.

### Construction of protein-protein interaction network

To assess the interconnectivity of DEGs in the RA synovium samples, we constructed a protein-protein network based on the interaction data obtained from public databases including BIOGRID (13), HPRD (14), IntAct (15), Reactome (16), and STRING (17). In the network, nodes and edges represent genes and functional or physical relationships between them, respectively. Graph theory concepts such as degree, closeness, and betweenness were employed to assess the topology of this network. Hub molecules were defined as the shared genes in top 10% with the highest rank in each arm of the three centrality parameters (18).

### Drug target prioritization strategy

To obtain a genome-wide drug target prioritization, we applied the Heat Kernel Diffusion Ranking approach (19). This method prioritizes the candidate genes by diffusing the differential expression values of the candidate genes through the network based on the confidence scores of the associations or interactions and is a powerful network-based machine learning approach to identify putative drug targets. The diffusion process is formulated by using a Laplacian exponential diffusion kernel and a score is computed by multiplying the differential expression values with the heat kernel. Drug targets are assumed to get a high score since these genes tend to be the central nodes in subnetworks showing significant transcriptional changes following treatment. The predictive power of the method is dependent on the network characteristics, the quality of the expression data, the type of drug and the target (19).

### Non-negative matrix factorization and determination of the optimal number of clusters

To classify the RA patients into subgroups based on their molecular signatures, we used the non-negative matrix factorization (NMF) method. NMF clustering is a powerful unsupervised approach to identify the disease subtype or patient subgroup and discover biologically meaningful molecular pattern (8,20). We applied the consensus NMF clustering method and initialized 100 times for each rank *k* (range from 2 to 6), where k was a presumed number of subtypes in the dataset. For each *k*, 100 matrix factorizations were used to classify each sample 100 times. The consensus matrix was used to assess how consistently sample-pairs cluster together. We then computed the cophenetic coefficients and silhouette scores for each *k*, to quantitatively assess global clustering robustness across the consensus matrix. The maximum peak of the cophenetic coefficient and silhouette score plots determined the optimal number of clusters (20). To confirm unsupervised clustering results, we used *t*-distributed stochastic neighborhood embedding (*t*-SNE) (21), a powerful dimensionality reduction method. The *t*-SNE method captures the variance in the data by attempting to preserve the distances between data points from high to low dimensions without any prior assumptions about the data distribution.

### Scoring of pathway activation

To quantify certain biological pathway activity, we calculated the gene expression z-scores (8,22). Briefly, a *Z*-score is defined as the difference between the error-weighted mean of the expression values of the genes in each pathway and the error-weighted mean of all genes in a sample after normalization. BCR-, chemokine-, Jack-STAT-, MAPK-, NFκB-, p53-, PI3K-AKT-, RIG-I-like receptor-, Fc ε RI-, TCR-, TGFβ -, TLR-, TNF-, VEGF-, and Wnt signaling pathways and their gene sets were imported from Kyoto Encyclopedia of Genes and Genomes (KEGG) database (23) and IFN type I- and type II signaling pathways and their gene sets referred to Reactome database (16). *Z*-scores were computed using each pathway in the signature collection for each of the samples, resulting in a matrix of pathway activation scores.

### Supervised learning analyses for the prediction of drug responsiveness

We used Naïve Bayes (NB), Decision Trees (DT), *k*-Nearest-Neighbors (KNN), and Support Vector Machines (SVM) to create drug responsiveness predictors (24,25). Each binary SVM was built using Gaussian Radial Basis Function (RBF) kernel and the Sigma hyperparameter was determined from the estimation based upon the 0.1 and 0.9 quantiles of the samples. For soft margins, the C parameter that achieved the best performance was in the range of 2^-4^ to 2^7^. For KNN, the *k* parameter was tuned in the range 2 to 20. All tuning hyperparameters were separately determined for each bootstrapped training dataset.

To determine the optimal feature set that enables distinguishing ‘good’ from ‘not good’ responders with the highest accuracy, we employed the wrapper feature selection method (25). The wrapper method uses the classifier as a black box to rank different subsets of the features according to their predictive power. In the wrapper method, a feature set is fed to the classifier and its performance is scored and the feature set with the highest rank is selected as the optimal feature set. The predictive power of each predictor was assessed through Receiver-Operator Characteristics (ROC) and Precision-Recall (PR) curve (26). Data was separated into independent training and test sets in a three-to-one sample-size ratio in a way of stratified random sampling. To make up for small sample size and minimize the error, we constructed the pool of resampled dataset by applying bootstrapping with 1000 iterations and subsequently applying a stratified 10-fold cross-validation (CV) for each bootstrapped dataset (24,25). Tenfold CV measures the prediction performance in a self-consistent way by systematically leaving out part of the dataset during the training process and testing against those left-out subset of samples. Compared to the test on independent dataset, CV has less bias and better predictive and generalization power. The predictive ability of the models generated from all the approaches was tested by performing the CV test at all the ten locations under study. Given the unequal numbers of trials in each class, balanced accuracy formula was employed to calculate the accuracy (27). The baseline is estimated by random expectation based on the pre-determined ratio of each condition. In case of IFXM, a probability of 0.29 (18/62) for a “good” and 0.71 (44/62) for a “not good” responder was applied.

### Statistical analysis

For continuous distributed data, between-group comparisons were performed using the one-way ANOVA, unpaired *t*-test or Mann-Whitney *U* test. Categorical or dichotomous variables were compared using the chi-squared test or Fisher’s exact test. To investigate the difference of pathway activation score across the subgroups, we fitted the one-way ANOVA model using logistic regression. All analyses were conducted in *R* (The R Project for Statistical Computing, www.r-project.org) and R packages used in the analysis and their references were summarized in the **Table S2**.

## Results

### The RA transcriptomics compendium

To get a list of RA-related DEGs, gene expression profiles of RA patients were compared with samples from the OA and NC groups. We identified 2762 DEGs for RA versus OA, and 3087 DEGs for RA versus NC (**Fig. 1**). Distribution of DEGs was assessed after the DEGs were divided into up- and down-regulated groups (**Fig. 2A**). The number of up-regulated DEGs was 1486 for RA versus OA and 1774 for RA versus NC. The intersection between two up-regulated DEG sets was 876, which we considered as RA-unique (**Fig. 2A** and **Supplementary File S1**).

**Figure 2.**
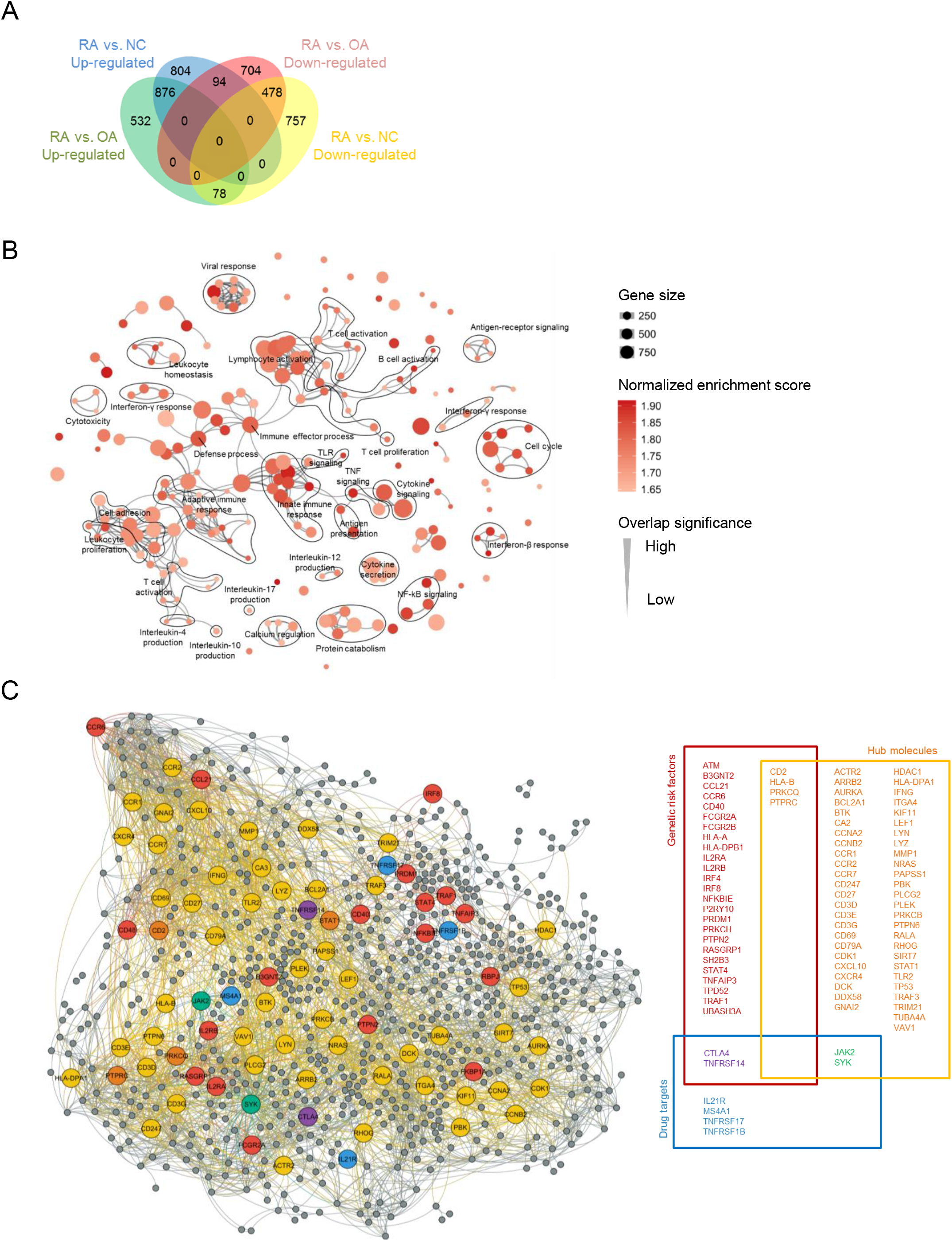
Differentially expressed genes and their functional network. **(A)** Venn diagram showing the overlap of up- and down-regulated DEG between RA versus OA and RA versus NC. **(B)** Gene-Set enrichment map for up-regulated DEG. Nodes represent GO-termed gene-sets. Their color intensity and size is proportional to the enrichment significance and the gene size, respectively. Edge thickness represents the degree of overlap between gene sets and only edges with a Jaccard coefficient larger than 0.25 were visualized. Clusters of functionally related gene-sets were manually curated based on the GO parent-child hierarchy and assigned a label. **(C)** Protein-Protein interaction network of up-regulated DEG. Red and blue nodes indicate the known RA-susceptible genes and drug target molecules, respectively. Drug targets were defined subject to the targets of drugs currently in use or under clinical trial and development. Yellow nodes correspond to the hub molecules, which are determined as the shared genes in top 10% with the highest rank in each arm of three centrality parameters; degree, closeness, and betweenness. Orange, green, and purple colored-nodes are the overlapped between red and yellow, yellow and blue, and red and blue ones, respectively. Right-side inset box is the schematic diagram of the interesting genes.

### Enriched biological processes and protein-to-protein interaction network

Through GSEA, we performed a functional enrichment analysis where 206 gene ontology processes were identified (**Fig. 2B** and **Fig. S3**). As expected, immune-related biological processes including adaptive and innate immune response, T cell- and B cell activation and response, and cytokine-related responses, were enriched. These occupied the main positions in the network and closely connected to each other. Among cytokine-related processes, interferon-β (IFN-β), interferon-γ (IFN-γ) interleukin (IL)-4, IL-10, IL-12, IL-17, toll-like receptor (TLR), and tumor necrosis factor (TNF)-related processes stood out as being substantially more enriched. Interestingly, several biological processes associated with viral invasion and defense response against viruses were newly identified (See **Fig. S4**). Metabolic processes such as calcium ion regulation and protein synthesis/transportation were enriched (all *P*<0.01), suggestive of active intracellular signaling and enhanced protein production and enzyme activity.

Identification of central attractors in the gene and protein network can provide targets for further experimentation and/or drug discovery. For this reason, we constructed the protein-to-protein interaction network of RA (**Fig. 2C**). We identified 3563 interactions among the 876 DEGs. Thirty-one of DEGs were overlapped with RA genetic susceptibility loci previously discovered (28) (**Fig. S5**) and a total of 56 genes were ranked as hub molecules based on the centrality analysis. The *CD2*, *PTPRC* (protein tyrosine phosphatase, receptor type C, also known as *CD45*), and *PRKCQ* (protein kinase C theta) were RA-susceptible genes having hub position in the network and products of these genes are involved in signal transduction of T cells. Eight genes including primary targets (*JAK2*, *SYK*, *CTLA4*, *MS4A1*) and counterpart receptor molecules (*TNFRSF14*, *TNFRSF17*, *TNFRSF18*, and *IL21R*) of cytokines targeted by the drugs currently in use or under clinical trial or development are also differentially expressed (29,30). Interestingly, the targets of small molecule therapeutics, *JAK2* and *SYK* are central hub nodes, in contrast to the targets of biologic agents, such as *CTLA4*, *MS4A1* (also known as *CD20*), *TNFRSF14*, *TNFRSF17*, and *TNFRSF18*. We found 219 RA-associated genes from the DisGeNet database (31), which are genes and variants having a responsible role in disease. Forty-six of them were overlapped with the RA synovial DEG. To assess topological proximity between RA-associated genes and drug targets in PPI network of synovial DEGs, the shortest distance between nodes was calculated (**Fig. S6**). Mean distance of *JAK2* and *SYK* was 2.11 ± 0.69 S.D. and 2.09 ± 0.68, respectively, and significantly shorter than those of other target molecules (range, 2.65 ~ 3.39) (in all cases p-value < 0.05).

### Identification and characterization of molecular subgroups

Next, we assessed whether RA patients can be categories in subgroups based on their expression profiles through consensus NMF clustering (20). To identify the optimal number of clusters and to assess robustness of the clustering result, we computed the cophenetic coefficient and silhouette score for different numbers of clusters from 2 to 6, where we found that 3 clusters are the optimal representation of the data (**Fig. 3A, Fig. S7**, and **Methods**). Segregation of RA subgroups was also reproduced by *t*-SNE analysis and principal component analysis (**Fig. 3B**). To identify characteristic molecular signaling pathways enriched in each cluster, we performed functional enrichment analyses for the predicted genes of each cluster (**Fig. 3C**). Nine enriched pathways were identified across the 3 clusters. RA cluster 1 (541 DEGs, 112 samples) exhibited activity for chemokine-, p53-, TNF- and TLR signaling pathways. Cluster 2 (130 DEGs, 64 samples) was also enriched in chemokine-, and p53 signaling pathways, while cluster 3 (205 DEGs, 80 samples) had a high enrichment of chemokine-, Fc ε RI-, Jak-STAT-, and T cell receptor (TCR) signaling pathways.

**Figure 3.**
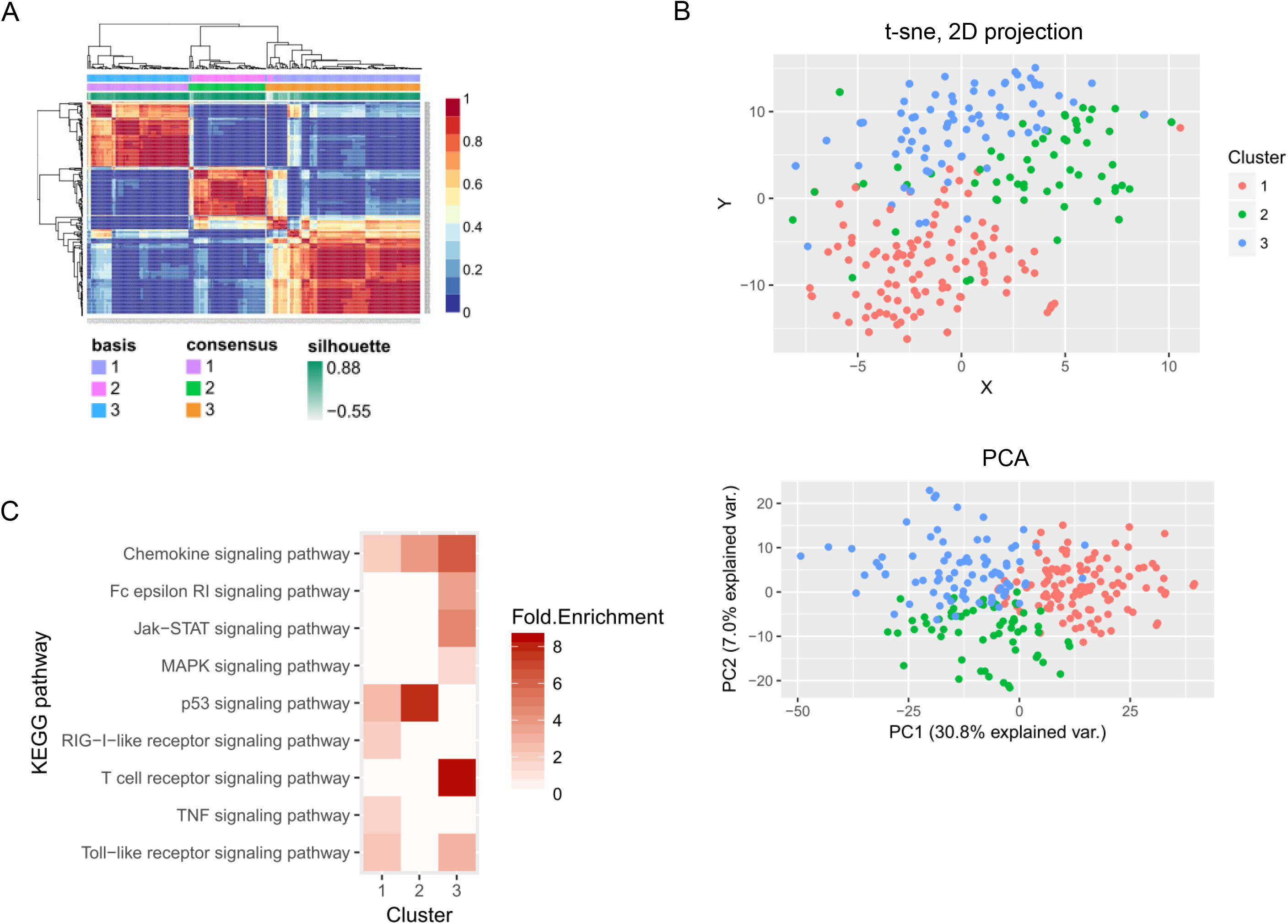
Identification of novel RA subgroups according to synovial signatures. **(A)** Reordered consensus matrices on RA compendium. The samples were clustered using average linkage and 1-correlation distances. Deep-red color indicates perfect agreement of the solution, whilst blue color indicates no agreement (Right-side color bar). Basis and consensus represent clusters based on the basis and consensus matrices, respectively. The silhouette score is a similarity measure within its own cluster compared to other clusters. **(B)** t-SNE (upper plot) and PCA (lower plot) reduces the dimensions of a multivariate dataset. Each data point is assigned a location in a two-dimensional map to illustrate potential clusters of neighboring samples, which contain similar gene expression patterns. **(C)** KEGG pathways enriched by the clustered DEG. Heatmap with red gradient represents the level of fold enrichment for each KEGG pathway, which was determined by DAVID software.

Some limitations lie with functional enrichment analysis using DAVID: dependency on the gene list not their expression levels, missing gene sets, and bias towards well-studied genes, and dropouts of some genes during combining the datasets. Thus, to better understand the differences among the three clusters, we analyzed the activation of individual pathways with adding six more pathways that are known to be associated with RA (30,32,33): transforming growth factor (TGF)- β-, vascular endothelial growth factor (VEGF)-, Wnt-, B cell receptor (BCR)-, NFκB-, PI3K-AKT (**Fig. 4**). As shown in the chord diagram, these pathways are strongly connected, with only TGFβ-, P53-, and Wnt signaling pathways more isolated than others (less shared DEGs). Especially TGFβ - and Wnt, have an opposite trend in their DEG expression (higher in cluster 1, mid in cluster 2 and low in cluster 3), which is the opposite of the trend we observe in most of the other pathways (**Fig. 4** and **Fig. S8**). P53 signaling pathways shared fewer genes with other pathways but strongly correlated with BCR-, chemokine-, Fc ε RI-, TCR-, TLR-, and TNF signaling pathways.

**Figure 4.**
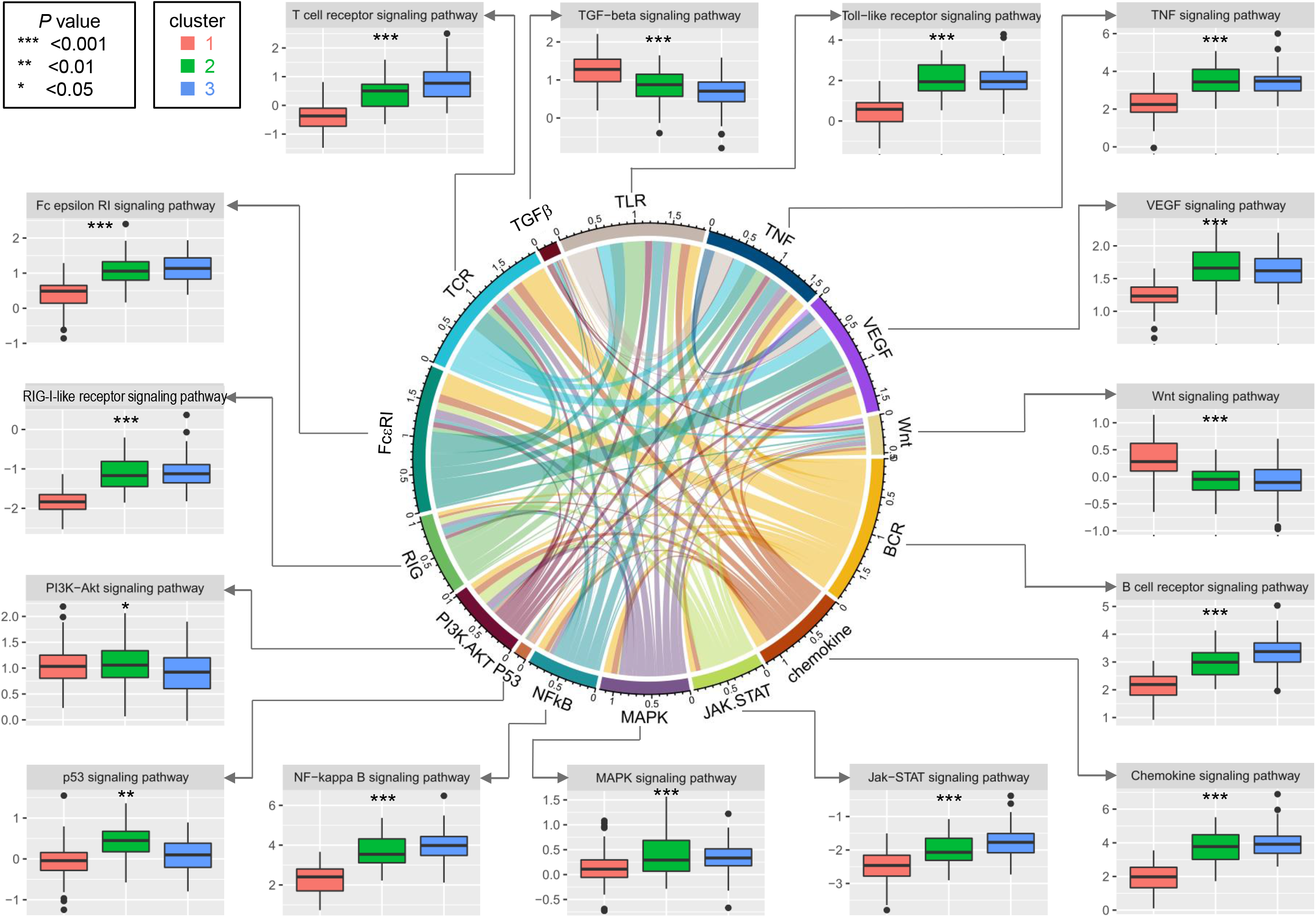
Pathway activation scores according to RA subgroups. Chord diagram shows interrelationship among pathways and link thickness is proportional to the overlap between two pathways, calculated using the Jaccard coefficients. Turkey boxplots reveals pathway activation scores across the RA subgroups and ANOVA test was used to analyze the differences among groups. *, *P*<0.05; **, *P*<0.01; ***, *P*<0.001.

While the activation scores of all pathways exhibited significant difference across the various clusters, all clusters exhibited one of the two trends in a statistically significant manner (*P*<0.05 in all cases) and in accordance with the observation through DEG-driven enrichment (all cases except TNF). It was found that IFN-related processes were enriched in GSEA and this was reproduced in the precedent studies (5,34-37). Since KEGG database did not provide IFN pathways data, we imported data on gene set of IFN pathways from the Reactome database. Levels of activation score of Type I and II IFN pathways were significantly different across the clusters (all *P*<0.001) and showed a tendency to increase from cluster 1 to 3 (**Fig. S9**). Compared with RA cluster 2 and 3, RA cluster 1 had moderate activation scores for most of the proinflammatory signaling pathways but high for PI3K-AKT-, TGF- and Wnt signaling pathways, which are principally involved in synovial proliferation and tissue remodeling (38). RA cluster 2 and 3 showed comparable activities for most of the proinflammatory pathways. More active in RA cluster 2 were the P53- and PI3K-AKT signaling pathways, which were reported to play a role in regulating apoptosis of synoviocytes or macrophages (39,40). In RA cluster 3, TCR-, Jak-STAT-, and NFκB signaling pathways were remarkable and it is noteworthy that IFN signaling pathways were most scored. Cellular processes affected by these pathways are in agreement with the DEG-driven enriched GO terms in each cluster (**Fig. S10**). This result indicates that there exist RA subgroups representing a distinct mode of inflammation deflected toward a certain combination of signaling pathways (**Table S3**). To prioritize the drug candidate targets, we ranked the DEGs using the Heat Kernel Diffusion Ranking approach (19). The identified drug target candidates are the central nodes in the subnetworks, with the highest disruption to the network under perturbation.

Next, we examined the relationship between identified 3 subgroups and the pertinent clinical features based on the provided information. There was no difference in gender ratio, age distribution, and tissue sampling method across the subgroups (*P* > 0.10 in all cases, see **Fig. S11**). Because data on the disease duration, activity, and RF positive were not provided individually for each sample, we compared two distinctively opposing datasets from compendium: the first (GSE45867) includes naïve, untreated RA patients with disease duration of <1 year, moderate disease activity and with arthroscopic needle biopsy performed before MTX or TCZM therapy (41). The second (GSE21537) is a cohort of the long-standing RA patients with high disease activity who had failed at least two DMARDs (including MTX) and did arthroscopic needle biopsy before IFXM therapy (42). Disease duration and activity were significantly longer and higher in the latter dataset (all *P* < 0.001) while there was no difference in age, gender, and RF positive between two datasets (all *P* > 0.10). Distribution of 3 subgroups did not differ between two datasets (*P* = 0.8664) (**Fig. S11**), indicating gene expression pattern by 3 subgroups have little direct relevance to disease duration and activity.

### Towards a predictor of drug response

For 105 RA samples that we had drug effectiveness data, we tested the hypothesis that there is an association between drug responsiveness and cluster membership. Out of the 5 drugs that we had data on (ADLM, IFXM, MTX, RTXM, and TOCM) we were not able to identify any such association (**Fig. S12**). Cluster 1 patients had an encouraging response to TOCM but at a low statistical significance level (p-value equal to 0.082). In addition to the intricacy of the pertinent pathways, the small size of samples treated by the specific drug, and their potential heterogeneity make the association between drug responsiveness and RA clusters difficult.

Since the differential expression of genes and pathways is at a higher resolution than general clustering signatures, we tested whether drug response can be predicted by using such features. We focused on the patients that were treated with IFXM due to the larger sample size (n=62). To test this hypothesis, we applied outcome to a binary classification (labels of “good” and “not good” responder) and tried two approaches: pathway-driven and DEG-driven models. Note that PCA analysis does not reveal separating distributions between the “good” and “not good” responders both for pathway activation score and DEG values (**Fig. S13**).

As features, we used the 17 pathways that are represented by continuous variables through their activation scores (refer to the pathway activation score for each pathway in the **Supplementary File S2**). To reduce the number of dimensions we performed feature selection through recursive elimination (**Table S4**). Based on those results made a predictive model using 4 supervised machine learning methods (NB, DT, KNN, and SVM) for selected key pathway scores and calculated the performance. All models outperformed the baseline (all *P* < 0.001) (**Fig. 5A**, left plot) and SVM, the best performing model, had an average performance AUC/AUPR of 0.87/0.78 (all *P* < 0.001) (**Fig. 5A**, middle and right plots). The selected key predictors for SVM model were NFκB-, Fcε RI-, TCR-, and TNF signaling pathways. Next, models based on expression values of DEG were fit in order to sort out the informative genes and compare their performance with pathway-driven models. DEG-driven models showed superior performance as compared with pathway-driven models (**Fig. 5B**, left plot). The overall AUC of the ROC curves exceeds 0.85 (**Fig. 5B**, middle and right plots). SVM showed the best performance AUC/AUPR of 0.92/0.86 and with the *HMMR*, *PRPF4B*, *EVI2A*, *RAB27A*, *MALT1*, *SNX6*, and *IFIH1* genes as features. The expression of these genes provide a distinct signature between two different outcomes (*P* < 0.05 in all cases, see **Fig. S14**).

**Figure 5.**
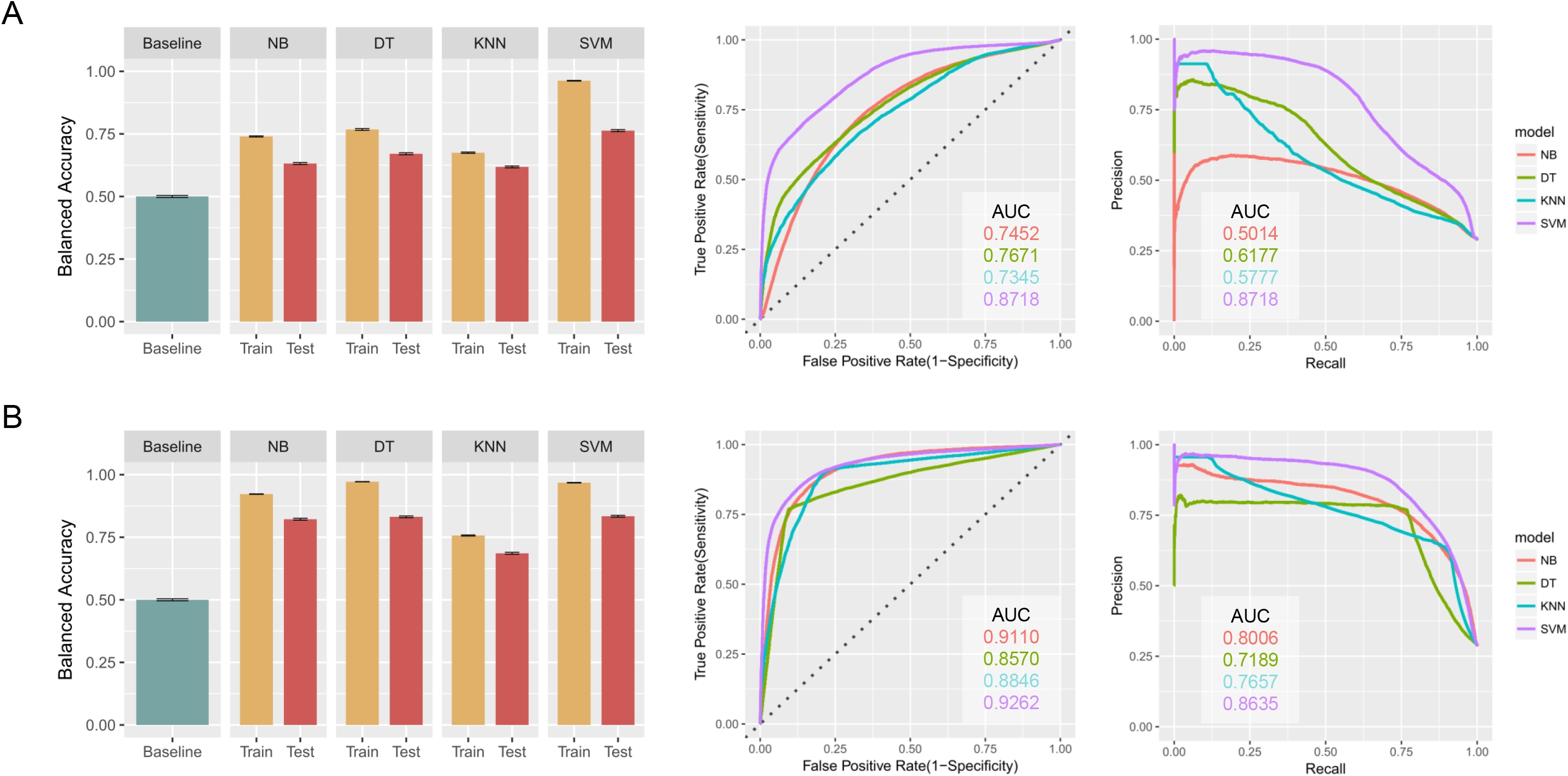
Predictive models and their performance. (A) Pathway-driven models.(B) DEG-driven models. (Left plot) The training and testing balanced accuracy for each classifier as compared with the baseline. All models outperformed the baseline (all *P* < 0.001) and the performance of the trained models was significantly compromised in testing sets (all *P* < 0.001). (Middle and right plots) Averaged ROC and PR curves showing the performance of each classifier.

## Discussion

Here, we built the largest RA compendium made by synovial transcriptomes. DEGs extracted from this compendium encompassed the susceptible genes and target molecules. Their topology in the network has opened new possibilities to elucidate biological roles and offer a cue for existing clinical questions. Unbiased cluster analysis of RA compendium resulted in meaningful categories of RA patients with distinct activity for relevant pathways. The pathway-based analysis allowed refinement in our understanding of RA subgroups and it was also feasible to construct pathway- or DEG-driven predictive model for intended treatment by machine learning methods.

Synovial tissues are considerably more difficult samples to obtain, as they are obtained during joint replacement surgery, synovectomy or by arthroscopy at 4-8 sites of the affected joint. However, they are more suitable to understand the mechanism and response to RA, since blood-derived samples are a distant and hence more noisy proxy to the disease, with known quality issues (4-6). Moreover, to refine the RA-unique genes, we compared RA samples with two control sets (OA and NC groups) and adopted the DEGs shared by three independent methods. We found that 24 of the DEGs are the known RA-associated genetic loci and take a central position in the synovial network. Since functional implications of risk allele were often obscure, it would be helpful to elucidate the biological mechanisms in which risk alleles operate. *STAT1*, a transcription factor downstream of IFN signaling pathway, highlighted as a key molecule in the previous reports (35,43), was found to be one of the hub genes. Other hub genes, such as *JAK2*, *SYK*, and *BTK* are small molecules that have increasingly drawn attention as novel therapeutic targets following the cytokine-targeting biologics (30). In contrast, molecules such as TNF receptor molecules, *CTLA4*, *IL6R*, and *MS4A1* were located at the functional periphery of the network although drugs against these molecules are widely used in clinical practice. Moreover, these molecules were placed farther from RA-associated genes than *JAK2* and *SYK* in the network, inferring part of their less potent efficacy in active RA. This was in good harmony with a recent clinical trial that baricitinib, an inhibitor of the Janus kinases *JAK1* and *JAK2*, showed a stronger therapeutic effect as compared with ADLM, a TNF inhibitor (44).

Biological processes and pathways identified from RA compendium show what is happening in the inflamed synovium of RA and are in good line with the previous studies (5,35,36). It is worthy of note that processes concerning viral cycle and anti-viral response were found to be enriched. This could be the internal process analogous to or the vestige of viral infection such as *Chikungunya* virus (45,46). A series of studies pointed out activation of IFN-related gene signatures in a subset of RA patients and its substantial similarity to viral infection (5,34-37,46) and one reported that the type I IFN signature negatively predicts the clinical response to rituximab treatment in patients with RA (34). Here, our results suggest that such a probable link between the IFN signature and the anti-viral response may exist (46).

Interestingly, we were able to identify three distinct subgroups through NMF analysis of the RA compendium and they differed in activation level of RA-relevant signaling pathways (8,20). Various combinations of molecular perturbations might converge to dysregulation of common pathways and lead to the similar phenotype (47). Since combinations of genomic perturbations are variable across the patients, pathway- or module-based approaches are desirable for a better understanding of complex inflammatory disease like RA. We looked at the enriched pathways derived from DEGs, which were commensurate with the pathway activation scores calculated from the whole gene list in the compendium. The RA cluster 1 was weighted toward signals regarding synovial proliferation and tissue remodeling (PI3K-AKT-, TGFβ- and Wnt signaling pathways) (38). RA clusters 2 and 3 showed a strong disposition for proinflammatory signaling pathways (Chemokine-, TNF-, TLR- and VEGF signaling pathways). Apoptosis-related pathways (P53- and PI3K-AKT signaling pathway) were much prominent in RA cluster 2 (39,40), while BCR-, Jak- STAT-, NFκB-, and TCR signaling pathways were stronger in RA cluster 3. It is known that synoviocytes are the main culprit of invasive synovium and quantitative and qualitative activities of synovial macrophage reflect therapeutic efficacy (48,49). They add to the cellular resistance to apoptosis and increase of the potential for proliferation, hence they contribute to the progression and perpetuation of destructive joint inflammation. Therefore, we speculate that an aggressive suppression of pro-inflammatory signals would be better pertinent to RA cluster 3, while therapeutic strategies to control propagation and survival of synoviocytes and macrophages together with anti-inflammatory treatment should be considered in RA cluster 1 and 2 (50) (**Table S3**). This insight, together with the candidate gene targets for drug development that we have identified in each cluster, may provide good starting points for delivering precision and personalized treatment.

Machine learning has become ubiquitous and indispensable for solving complex problems in most sciences (51). Since the problem of unresolved heterogeneity is prevalent to medicine, the same methods are expected to open up vast new possibilities in medicine and actively employed in a variety of clinical research (51). We tried to make a predictive model for 62 samples that were obtained from the synovial tissue of RA patients before administration of IFXM. Because key features are informative for predicting the outcome rather than being directly implicated in the major pathways or usual suspects related to the RA synovium, they could be different depending on drugs and models. The fact that we achieved high performance scores in RA response prediction from mining the RA compendium, despite this was not attainable through individual statistical techniques in the past (42), argues that similar techniques can guide us to narrow choices for more effective drugs. Interestingly, DEG-driven models outperformed models that were relying on pathways as features. Among 7 featured genes in SVM model, HMMR (Hyaluronan-mediated motility receptor, also known as RHAMM) exacerbated collagen-induced arthritis by supporting cell migration and up-regulating genes involved with inflammation (52) and MATL1 (Mucosa associated lymphoid tissue lymphoma translocation gene 1) was recently identified to play a crucial role in the pathogenesis of RA as MATL1-deficient mice were completely resistant to collage-induced arthritis (53). Direct connection to RA was not revealed for the rest of the identified informative genes so far and it remains to be investigated how and why these features are indicative of drug response.

There are some limitations to be addressed in this study. First, removal of batch effects is not ideal which adds to the noise in the compendium. Second, we did not fully address the association of RA subgroup with clinical factors including age, sex, disease duration, and RF positive due to lack of complete annotation for each RA sample. Third, a limited number of samples were treated with other drugs except for IFXM precluded us from making a predictive model. In general, more meta-data would be desired, although this is to be expected as these studies were performed in different clinical environments, with different procedures and goals, which did not include their aggregation to a single compendium and application of advanced machine learning techniques. In the future, we anticipate that the construction of datasets with sufficient metadata for machine learning analysis would enable critical insights and may lead to novel drug targets for RA treatment.

## Author Contributions

K-J Kim, I Tagkopoulos designed the study and K-J Kim acquired the data. K-J Kim, M Kim, I Tagkopoulos contributed to the analysis and interpretation of data. K-J Kim conducted all data analysis and drafted the initial manuscript. All authors were involved in drafting the article or revising it critically for important intellectual content, and all authors approved the final version to be published.

